# Assessing large-scale genomic language models in predicting personal gene expression: promises and limitations

**DOI:** 10.1101/2025.07.09.664024

**Authors:** Shumin Li, Ruibang Luo, Yuanhua Huang

## Abstract

Large-scale genomic language models (gLMs) hold promise for modeling gene regulation, yet their ability to predict personal gene expression remains largely unexplored. We developed a framework, gLM2X-Tower, to benchmark gLMs and sequence-to-function (S2F) models on this task with paired personal genome-transcriptome data. With individual-level training, we found that similar to S2F models (e.g., AlphaGenome), gLMs (e.g., Evo2) remain incapable of predicting the inter-person variability on held-out genes. However, such training improves prediction for seen genes in new individuals, particularly by gLMs, highlighting the potential applications in few-shot settings like for rare variants

## Background

Building accurate genotype-to-phenotype models remains central to modern biology, often with genotype-to-expression as the junction. While expression quantitative trait loci (eQTL) studies have identified thousands of loci associated with gene expression, these approaches are primarily limited to common variants due to their reliance on population-scale data. As a result, rare and novel variants—key to understanding disease mechanisms and regulatory biology—remain difficult to interpret.

Sequence-to-function (S2F) models such as Enformer [1] and Borzoi [2] address this gap by mapping genomic sequence to regulatory outputs using deep learning. Trained on reference genomes with multiomic labels across diverse cellular contexts, they accurately predict average gene expression across genes and cell types. However, their ability to capture the transcriptomic variation across individuals, driven by personal genetic variants, remains limited [3, 4]. While AlphaGenome was recently released to improve variant effect prediction, its ability to model gene expression from personal genomes also has not yet been evaluated [5].

Genomic language models (gLMs) offer a complementary approach. Pretrained via self-supervised learning on DNA sequences, they learn generalizable representations for downstream tasks and showed promising results [6, 7]. Yet their ability to model individual gene expression—and whether they can overcome the limitations of S2F models—has not been evaluated.

On the other hand, S2F models and gLMs share structural similarities, including self-attention-based encoders and task-specific heads. However, they diverge in training strategy and data. S2F models are trained with multiomic labels across diverse cellular contexts, enabling implicit contrastive learning of regulatory grammars. In contrast, gLMs are pretrained solely on DNA sequence data and fine-tuned with unimodal outputs. Here, two questions remained unexplored: (1) How much can model performance improve by increasing genetic diversity in the training data (for example through more individual genomes) before hitting a ceiling that only new modeling techniques can break? (2) To what extent can gLMs trained solely on paired DNA–RNA data—without epigenomic annotations or multi-cellular labels—accurately predict individual-level gene expression in unseen genomic sequences?

## Results and discussion

To address the questions, we benchmarked three gLMs—Evo2-7B [7], Nucleotide Transformer v2 (NT) [6] and Caduceus [8]—and three S2F models—Enformer, Borzoi and AlphaGenome—on their ability to predict personalized gene expression using paired genome–transcriptome data from GEUVADIS [9]. These data include lymphoblastoid cell line (LCL) RNA-seq and high-quality genotypes from 1000 Genomes Project (1KG) individuals. We used log-TPM processed by [10] and reconstructed personalized DNA sequences by substituting SNPs from phased 1KG genotypes into the reference genome. From a total of 3,259 genes with significant cis-eQTLs in European individuals, we selected 2,976 genes for training with reference sequence and population-average expression, 200 randomly sampled genes for training with personal genomes, and 287 genes on chromosomes 5 and 10 for held-out evaluation. The dataset included 50 individuals for training and 100 for testing. (Fig. 1A–B, Methods).

**Fig. 1.**
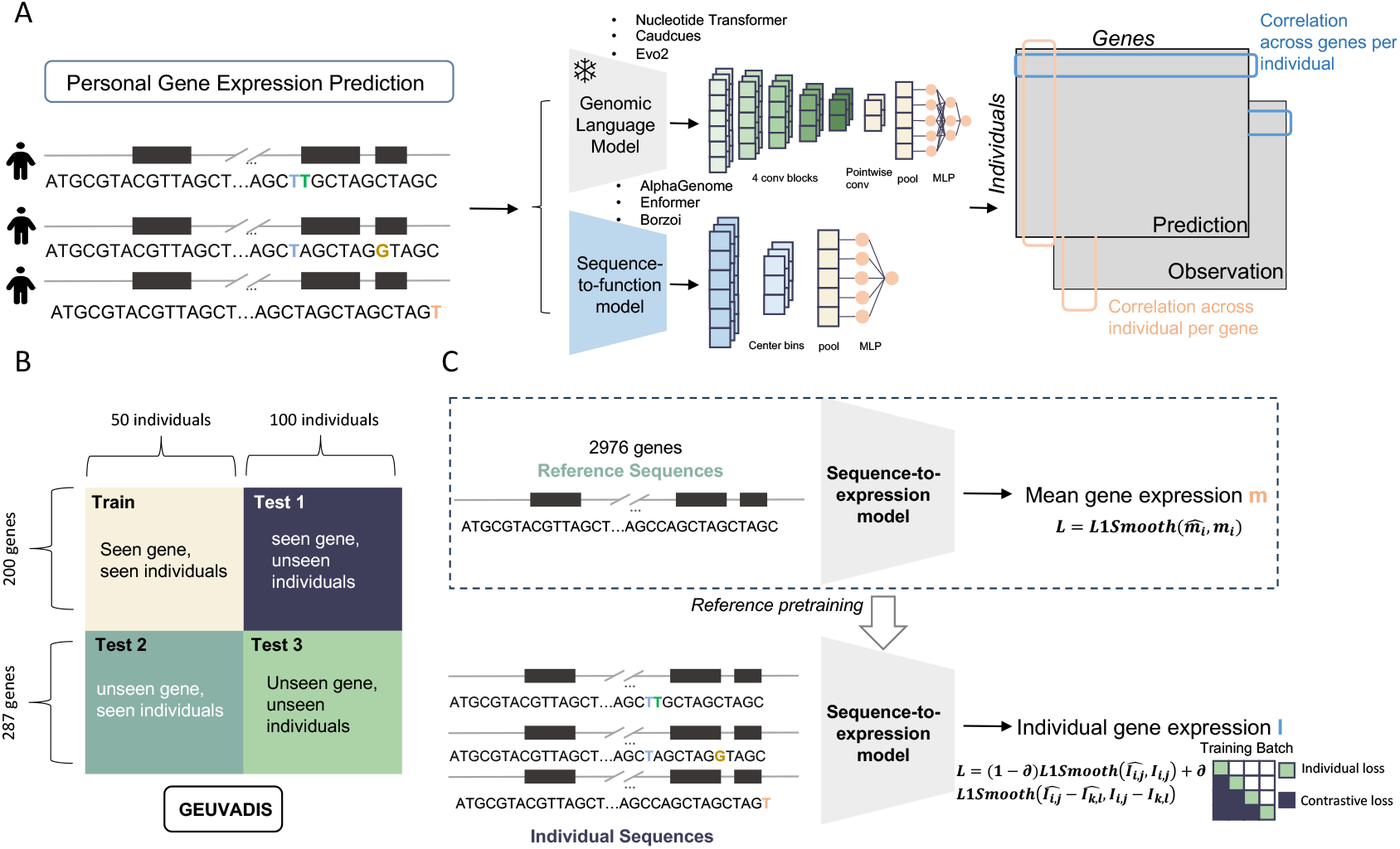
Benchmarking gLMs and S2F models for personalized gene expression prediction. A, Overview of this study: trained on matched genome-expression data, S2F models (Enformer, Borzoi) are fine-tuned by replacing with a task-specific head and embeddings from gLMs (Nucleotide Transformer, Caduceus, Evo2) are utilized for training a multi-layer CNN prediction tower. Noted that AlphaGenome was only evaluated on its original weights without fine-tuning. Model performance was evaluated based on both across-individual-and across-gene Spearman correlations. B, Individual-related data partitions across genes and individuals for the dataset of Geuvadis consoritum. C, Training strategies. Models were trained using one of three approaches: reference-only, individual-only, or reference pretraining followed by individual finetuning. Strategies involving individual genomes were optimized with a combination of individual-level and contrastive loss.

To fully utilize both gLM and S2F models, we developed “gLM2X-tower” a fine-tuning framework with an expression prediction tower (Fig. 1A). Specifically, we implemented three training strategies (Fig. 1C): (1) reference-only models trained on population-average expression and the reference genome (“r-”); (2) individual-only models trained on personalized sequences and matched expression profiles (“p-“); and (3) combined models pretrained on the reference genome and fine-tuned with individual-level data (“rp-”). For individual-based models, we applied a hybrid loss function that combined a smooth L1 loss on individual predictions with a contrastive component to emphasize inter-individual and inter-gene variations within each batch. Sepcifically, AlphaGenome was evaluated using its original model without fine-tuning, as its pretrained weights are not publicly available.

For the gene-by-individual expression prediction, there are two levels of tasks (Fig. 1A): cross-gene variability (row vector) that has been nearly solved by S2F models (also see our replications in Fig. 2C), and cross-individual variability (column vector), an unsolved challenge that we primarily focus here using seen and unseen genes (Tests 1 and 3, Fig. 1B). First, for seen genes, rp-Caduceus achieved performance comparable to the PrediXcan-style gene-specific linear model (Fig. 2A). Reference-only models failed to capture personal variation, while fine-tuning with individual genomes consistently improved correlation. For example, in the case of METTL18, the original Enformer showed a negative correlation of -0.247, while all fine-tuned models achieved Spearman correlations above 0.3 and AlphaGenome achieved 0.464 (Fig. S1). Similar improvements were observed for NBPF (Fig. S2): personal genome fine-tuning increased the correlation from 0.463 to 0.582 for Enformer and rp-Evo2 pushed it to 0.69. An exception was ZNF324B, where fine-tuning slightly reduced performance but maintained the correct direction (Fig. S3). Overall, AlphaGenome and Borzoi outperformed Enformer, likely due to its finer single-base and 32 bp compared to 128 bp resolution in embedding, and gLM generally outperforms S2F models when access to individual fine-tuning data (mean: 0.197 by rp-Caduceus vs 0.112 by rp-Borzoi). The case of HLA-DQA1 (Fig. S4) emphasizes this effect, suggesting that self-supervised training enhances generalizability. Notably, 46 genes that PrediXcan could not predict can be modeled by rp-Caduceus (Fig. 2B), showing the benefits of using the sequence-based models to increase the range of applicability, despite requiring knowing the variants and their linked genes.

**Fig. 2.**
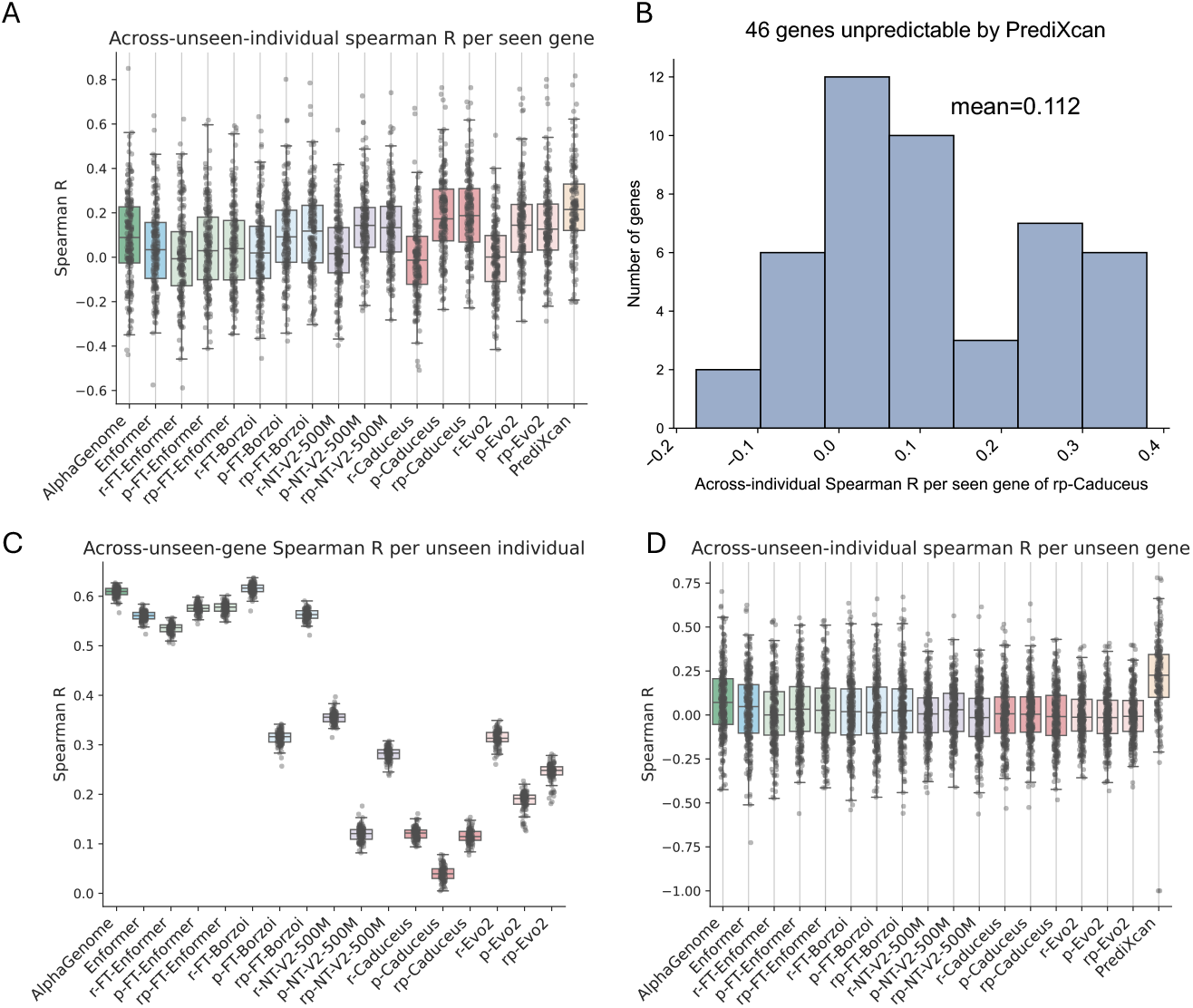
Comparison of personalized expression prediction across models and tasks. A, Across-unseen-individual correlation per seen gene for gLMs and S2F models (Test 1). The statistical model PrediXcan was trained per gene using genotypes of common variants (allele frequency 0.05). B, Correlation performance of *rp-Caduceus* on 46 genes poorly predicted by PredXcan. C, Across-unseen-gene correlation per unseen individual (Test 3). D, Across-unseen-individual correlation per unseen gene (Test 3). PredXcan was trained specifically on these unseen genes and serves as an approximate upper bound for predictive performance in this setting.

Next, we tested generalization to unseen genes in unseen individuals (Fig.2D). None of the models performed reliably in this setting. Although some showed a long-tailed distribution of predictive performance (suggesting a subset of genes may still be predictable, Fig. S5), overall performance remained low. These results show that current models—whether S2F or gLM-based—struggle to generalize across loci and individuals. This situation is not relieved even by incorporating more training individuals (Fig. S6).

Taking each gene as an independent output variable, we may view Test 1 as few-shot learning and Test 3 as zero-shot learning. Despite the limited power in a zero-shot setting, current models started to show promise in few-shot learning (Fig.2A), making it closer to applications with rare or somatic variants, especially if one may further reduce the required number of fine-tuning individuals. Moreover, increasing the number of training individuals leads to improved performance on Test 1, but not on across-unseen-individual variations on Test 3. This indicates a current bottleneck in the ability of sequence-based models to generalize across genomic loci.

## Conclusions

In summary, our results support three conclusions. First, incorporating personal genomes improves prediction for seen genes, but this effect does not generalize to unseen loci—highlighting the challenge of learning transferable *cis*-regulatory grammar. Second, multi-species pretraining (Evo2 and NT) outperformed human-only models like Caduceus in unseen contexts for population-average level prediction, emphasizing the benefit of evolutionary diversity. Meanwhile, S2F models trained with multi-omic data perform substantially better in this task (Fig. 2C), suggesting that such annotations provide important information. Third, despite the promise, gLMs do not overcome the current bottleneck in personal gene expression prediction for unseen contexts, even if the number of training individuals is increased (Fig. S6). Similarly, AlphaGenome—the most advanced S2F model optimized for variant effect prediction—also fails to generalize in this setting, although it performs better than its precursor, Enformer. Future work should focus on augmented training data—spanning both individual-level and multi-cellular contexts—along with efficient model architectures, improved training strategies, and a sharper application focus on causal variant identification. Several successful approaches from related sequence modeling tasks may be adaptable to this setting (Table S1).

## Methods

All data used in this study are publicly available (‘Availability of data and materials’) and no specific ethics approval was needed to conduct the current study.

### Personal gene expression dataset

We used paired whole-genome sequencing (WGS) and gene expression data from the GEUVADIS consortium [9], comprising lymphoblastoid cell lines (LCLs) from individuals in the 1000 Genomes Project [11]. A total of 150 individuals were randomly selected and partitioned into 50 for training and 100 for testing. This study focused on 3,259 genes identified in the original GEUVADIS study as having significant cis-eQTLs in European individuals. Of these, 200 genes—excluding those located on chromosomes 5 and 10—were randomly selected for training with individual sequences, while 2,976 genes were used for training with reference sequences. The remaining 287 genes from chromosomes 5 and 10 were reserved for testing. We used log-transformed TPM gene expression values and annotated transcription start sites (TSS) as processed by [10] and [3]. Additionally, we constructed a reference genome–based dataset using hs37d5 paired with population-average gene expression levels computed from all 462 individuals. For each gene, a 64,000 bp sequence window centered on the TSS was used as input to all models, except for Nucleotide Transformer (NT), which used 11,000 bp due to model design constraints and AlphaGenome, which used 131,072 bp selected from a list of supported input lengths. Sequences were reverse-complemented for genes located on the negative strand.

Individualized input sequences were constructed using phased genotypes from the 1000 Genomes Project. We generated personalized DNA sequences using *bcftools consensus* [12], including only singlenucleotide variants (SNVs) with the options -H 2pIu[1pIu] and -i “type=‘snp’”.

### Predicting gene expression with S2F models

#### Predicting gene expression with the original Enformer

We used the pretrained Enformer model available at [13]. For each input sequence, we extracted the predicted track corresponding to CAGE:epitheloid carcinoma cell line:HelaS3 ENCODE, biol. Predictions were averaged across the central 10 output bins. Both haplotypes of each individual were input separately, and their mean was used as the final gene expression prediction.

#### Predicting gene expression with AlphaGenome

We evaluated AlphaGenome via the publicly available Google API. Given its support for fixed input lengths (2,048 to 1,048,576 bp), we selected 131,072 bp to enable a comparison with other models, which used 64,000bp inputs. Predictions were generated by specifying the ontology term “EFO:0002784”, corresponding to the same LCL cell line used in our benchmarking dataset. For each input sequence, we averaged the output across the central 1,280 positions of the forward-strand CAGE track outputs, similar to the center-bin averaging used for Enformer. Both haplotypes for each individual were input separately, and the final gene expression prediction was computed as their mean.

#### Fine-tuning S2F models

We fine-tuned the full Enformer and Borzoi models by replacing their original task-specific heads with a regression head. Backbone parameters were initialized using pretrained weights from [1, 2, 13, 14]. The new head included attention pooling applied to the central 10 output bins, yielding fixed-size embeddings of 3,072 dimensions for Enformer and 1,920 dimensions for Borzoi. These embeddings were passed through a fully connected linear layer to generate scalar predictions. Final gene expression values were obtained by averaging predictions across both haplotypes of each individual.

For the reference-only and individual-only training strategies, models were optimized using the Adam optimizer [15] with an initial learning rate of 1e-6. A linear warm-up schedule was applied over the first 10% of training epochs, with a start factor of 0.01 and an end factor of 1.0, followed by a cosine scheduler for the remaining epochs. A total of 40 training epochs were used; no validation set was held out, and models from the final epoch were used for evaluation. For the combined reference and individual training strategy, models were first trained on reference-only data were utilized for fine-tuning with individual genome-expression pairs for 40 epochs with the same configuration. Experiments were conducted on NVIDIA H800 or L40S GPUs using a batch size of 8 and a random sampling strategy.

For reference-only training, we used Smooth L1 loss to optimize the models, where for individual-based training (including *p-* and *rp-* models), we defined a hybrid loss combining individual prediction accuracy with inter-sample consistency. The total loss ℒ is computed as:

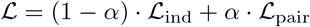

where:

- ℒ_ind_ is the smooth L1 loss between predicted and true gene expression values for each individual,
- ℒ_pair_ is a contrastive component that encourages the predicted pairwise differences to match the ground-truth differences across individuals and across genes,
- *α ∈* [0, 1] is a weighting factor (set to 0.3 in this study).

Let *ŷ_i_* and *y_i_* be the predicted and true expression values for individual *i*, respectively. The individual loss is defined as:

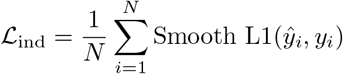

The pairwise contrastive loss is:

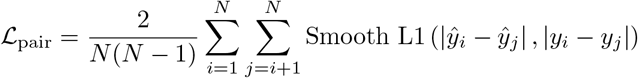

where *N* is the batch size, and Smooth L1(*·, ·*) denotes the Smooth L1 loss between two scalars.

#### Predicting gene expression with gLM embeddings

For gLM-based personal gene expression prediction, we used the output of layer blocks.28.mlp.l3 from Evo2-7B, the final layer from Caduceus-ph, and layer 20 from the NT as model embeddings. We constructed a multi-layer CNN to process the three gLM embedding inputs. The input to the model consists of a sequence of embedding vectors with shape (sequence length × embedding dimension), where the sequence length is 64,000 for Evo2-7B and Caudcues, 11,000 for NT, and the embedding dimensions are 4,096 for Evo2-7B, 256 for Caduceus-ph, and 1,024 for NT

The model begins with a stack of four blocks, each implemented as a depthwise separable convolutional module. Each block consists of:

- A depthwise convolution followed by batch normalization and ReLU activation,
- A pointwise (1×1) convolution followed by batch normalization and ReLU activation
- An optional attention pooling.

These blocks progressively reduce the sequence length from 64,000 to approximately 63 tokens. The block parameters (kernel size, stride, padding, and pooling size) are:

- Block 1: (16, 8, 7, 2),
- Block 2: (8, 4, 3, 2),
- Block 3: (4, 2, 1, 2),
- Block 4: (3, 2, 1, 1; no pooling).

Following the convolutional blocks, a pointwise convolution projects the embedding dimension to 512. The tensor is then processed by a global average pooling layer (AdaptiveAvgPool1d(1)), reducing it to a fixed-length vector of 512 features per sample.

The final classification head consists of two fully connected layers with ReLU activation and dropout (0.5), producing a scalar corresponding to predicted gene expression. And the model processes both haplotypes independently and averages their predictions to produce a final output.

For the training strategies of gLM-based models, we took the same training settings and the losses as described in **Fine-tuning S2F models**, except we set the initial learning rate as 1e-4.

#### Predicting gene expression with PrediXcan

We implemented a PrediXcan-style baseline using the ElasticNetCV model from scikit-learn, with 10-fold cross-validation, a fixed random state of 0, and all other parameters set to default. Separate models were trained for each gene using the same training and testing individuals as the sequence-based models, covering both 200 randomly selected training genes (results in Fig. 2A) and 287 held-out test genes (results in Fig. 2D). For each gene, we defined a 64kb window centered on the TSS and extracted individual genotypes within this region using a minor allele frequency (MAF) threshold of 0.05.

## Supplementary information

If your article has accompanying supplementary file/s please state so here.

Authors reporting data from electrophoretic gels and blots should supply the full unprocessed scans for key as part of their Supplementary information. This may be requested by the editorial team/s if it is missing.

Please refer to Journal-level guidance for any specific requirements.

## Declarations

### Ethics approval and consent to participate

Not applicable

### Consent for publication

Not applicable

### Competing interests

The authors declare no competing interests.

### Funding

### Authors’ contributions

Y.H. conceived and supervised this study with the support of R.L. S.L. implemented the gLM2X-Tower and performed all benchmarking analyses. S.L. and Y.H. wrote the manuscript, and all authors approved it.

## Acknowledgements

We would like to thank Dr. Ruiyan Hou for data exploration and Dr. Jiecong Lin for discussing sequence models.

## Availability of data and materials

Source code of this study is available at: https://github.com/StatBiomed/gLM2X-Tower; The paired genotype and RNA-seq data is obtained from Geuvadis consortium (https://www.ebi.ac.uk/biostudies/arrayexpress/studies/E-GEUV-1#processed-data) and the processed TPM value is obtained from https://github.com/ni-lab/finetuning-enformer/tree/main/process_geuvadis_data.

## Notes

### Competing Interest Statement

The authors have declared no competing interest.

