## Supplementary Information for "Assessing large-scale genomic language models in predicting personal gene expression: promises and limitations"

#### Supplementary Figures

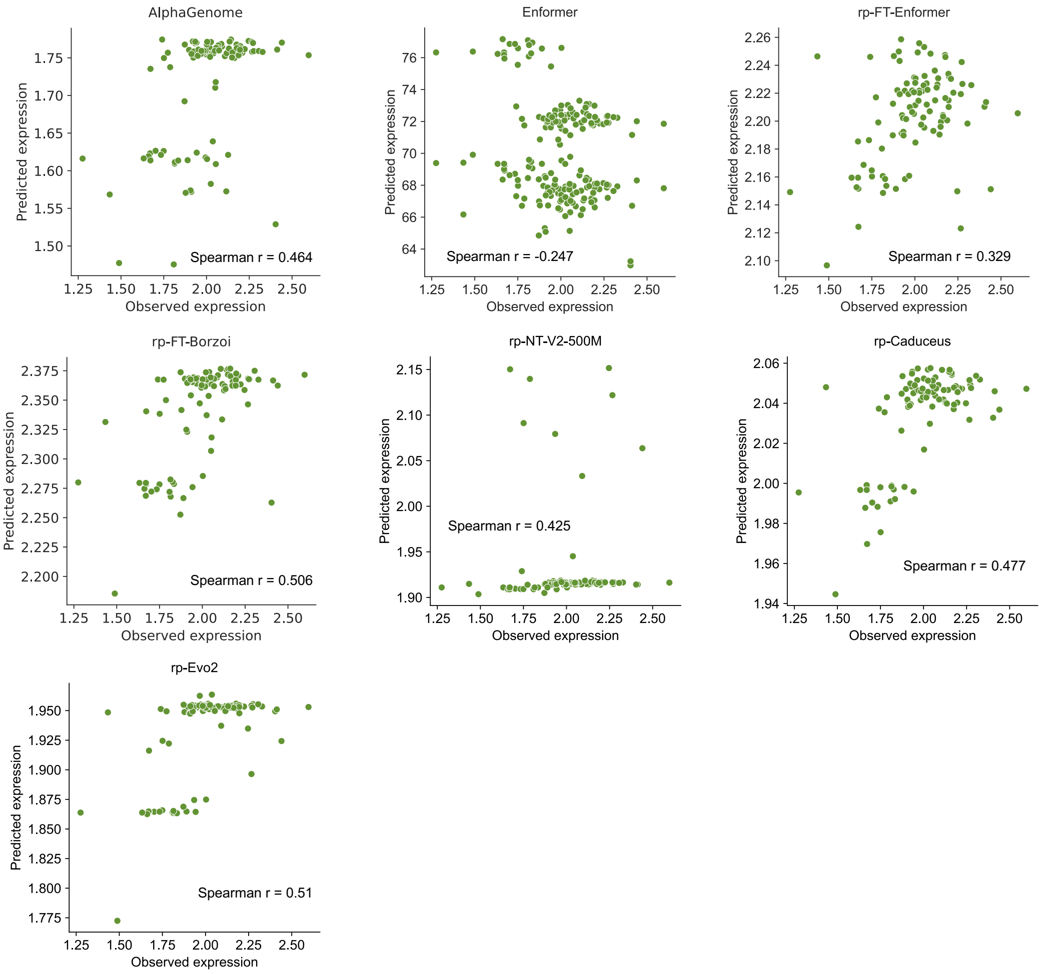


**Fig. S1  Model performance for METTL18.**

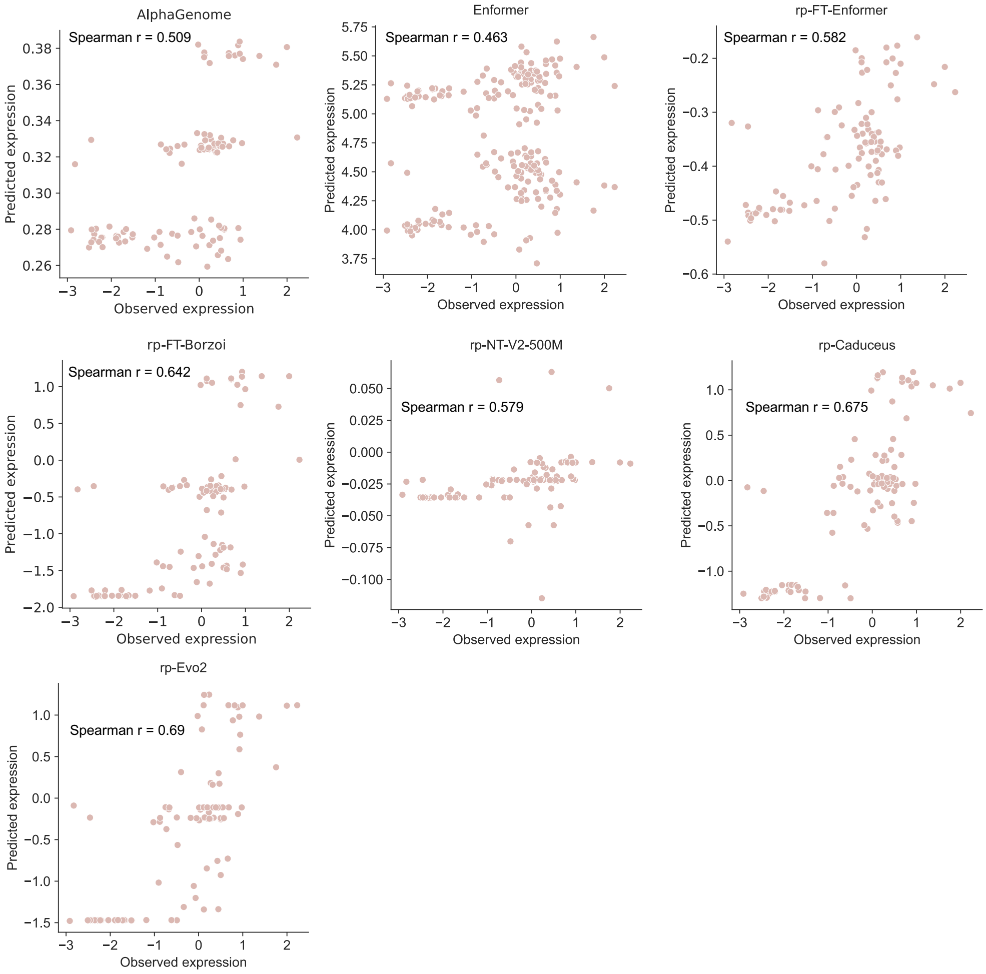


**Fig. S2 Model performance for NBPF.**

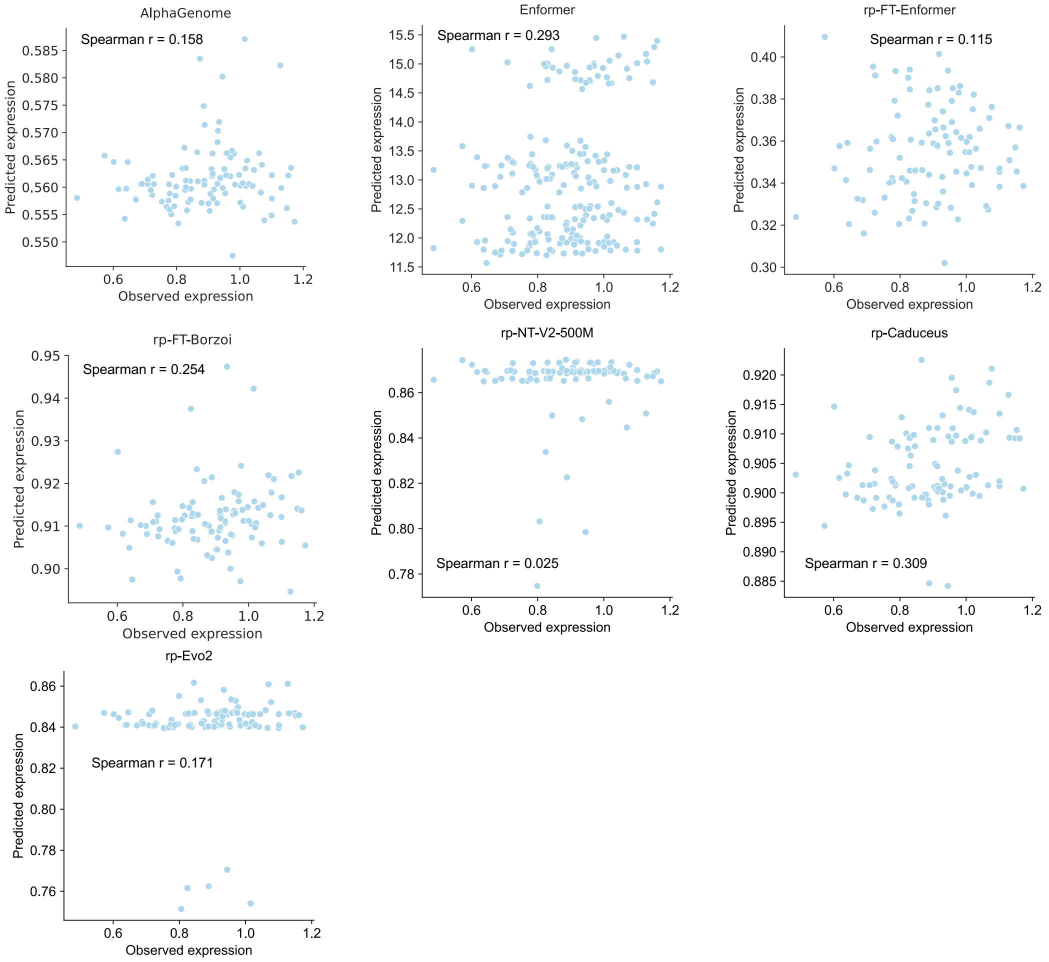


**Fig. S3** **Model performance for ZNF324B.**

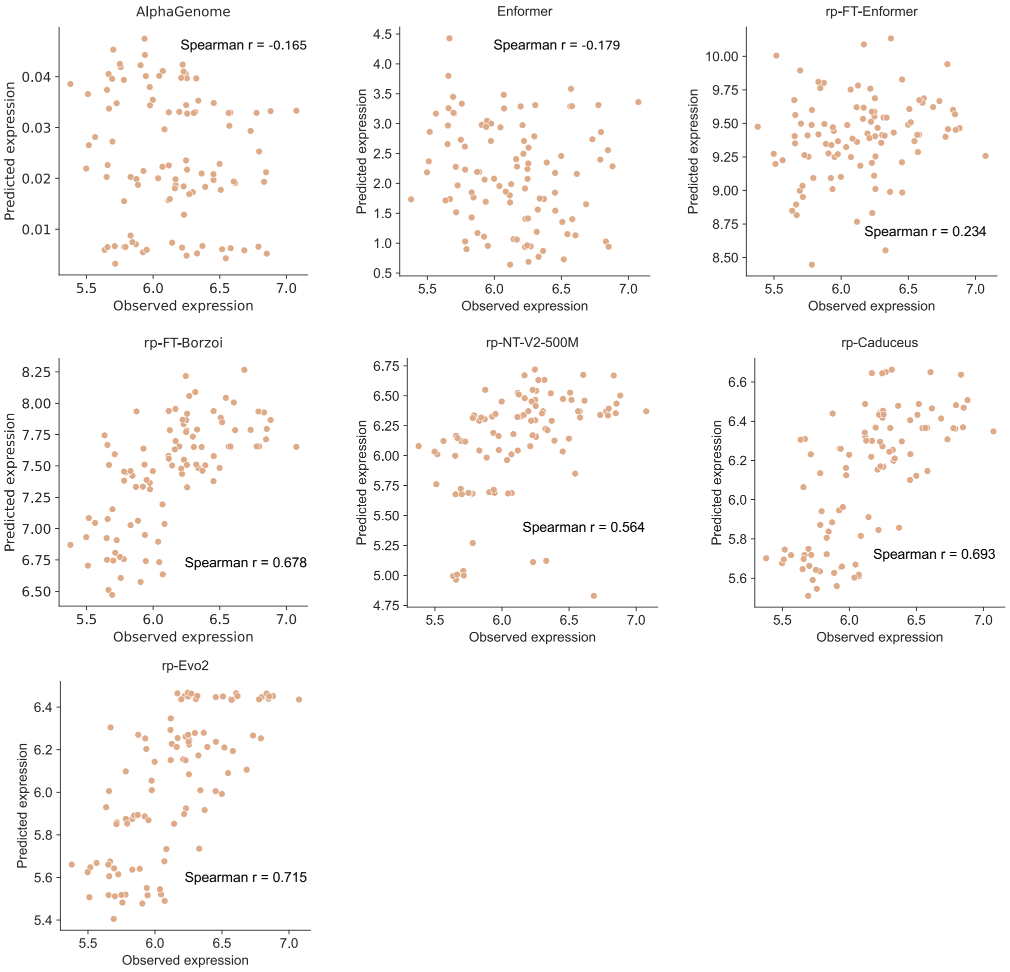


**Fig. S4 Model performance for HLA-DQA1.**

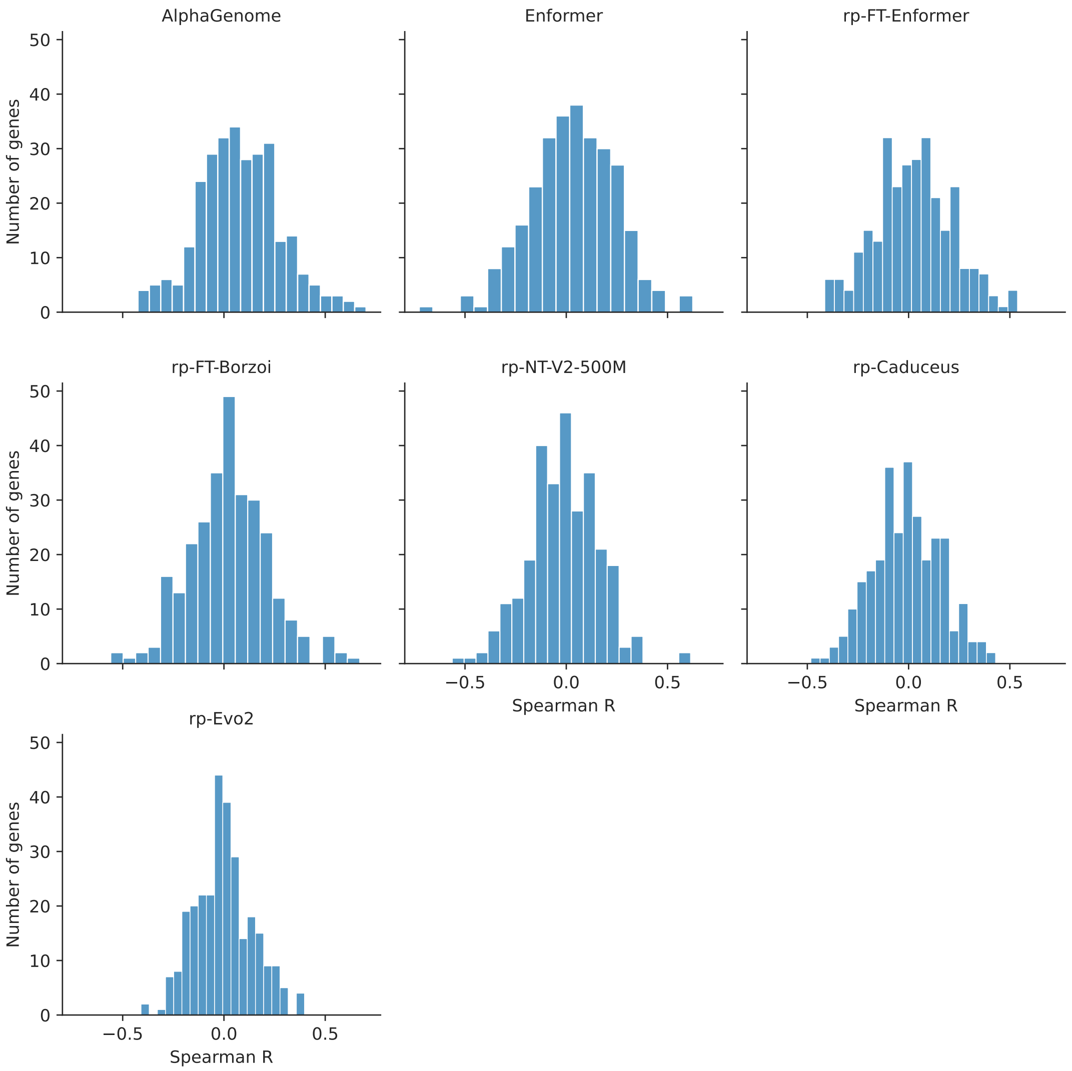


**Fig. S5 Distribution of Spearman R for unseen genes across unseen genes (task 3).**

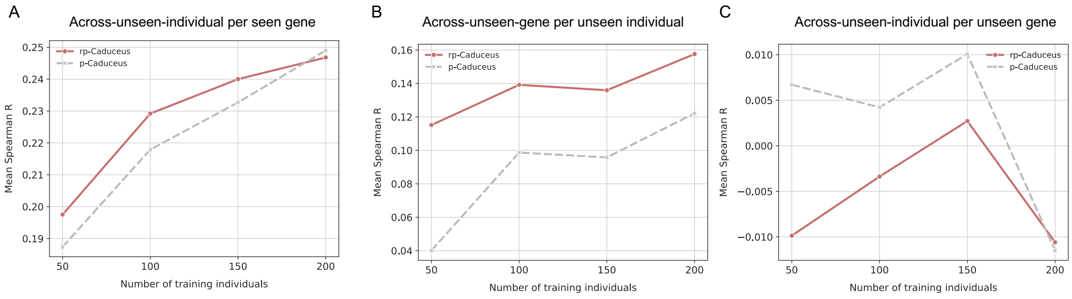


**Fig. S6 Performance of Caduceus-based models based on increased training individuals.** (A), Across-unseen-individual correlation per seen gene. (B), Across-unseen-individual correlation per unseen gene. (C), Across-unseen-individual correlation per unseen gene

### Supplementary Tables

**Table S1** Future directions for improving personalized gene expression prediction.

| Direction | Strategies | Examples |
|---|---|---|
| **Input Data** | Extend sequence length for distal elements | Borzoi (524kb) [1]; Evo2 (1M) [2] |
| | Increase number of training individuals | Large training set [3, 4] |
| | Incorporate epigenomics data | GET [5], EPInformer [6] |
| | Integrate multi-cellular labels | Basenji2 [7], Enformer [8], Borzoi [1] |
| | Utilize population LD data | SuSie [9] |
| **Model Architecture** | Use finer tokenization | Single-nucleotide resolution [2, 10] |
| | Scale model capacity | Evo2-70B |
| | Develop lightweight architectures | EvoFormer in AlphaFold3 [11] |
| | Improve training efficiency | Contrastive learning model architecture [12] |
| **Application Focus** | Prioritize causal variant | AlphaMissence [13], GPN [14] |
| | Fine-mapping and clinical interpretation | SuSie [9] |
